# Chitosan-Polyphosphate Scaffold Loaded with Copper for Endodontic Regeneration: A Laboratory Study

**DOI:** 10.1101/2025.10.30.685681

**Authors:** Hanan Moussa, Isabel Mello, Brendan Leung, Mark Filiaggi

**Affiliations:** Department of Biomaterials & Applied Oral Sciences, Faculty of Dentistry, Dalhousie University, Halifax, Canada; Department of Prosthodontics, Faculty of Dentistry, Benghazi University, Benghazi, Libya; Department of Prosthodontics, Faculty of Dentistry, Hespereds University, Benghazi, Libya; Department of Dental Clinical Sciences, Faculty of Dentistry, Dalhousie University, Halifax, Canada; School of Biomedical Engineering, Dalhousie University, Halifax, Canada; Department of Microbiology and Immunology, Faculty of Medicine, Dalhousie University, Halifax, Canada

**Keywords:** Regenerative endodontics, Chitosan, Polyphosphate, Copper, Stem cell

## Abstract

**Objective:** Regenerative endodontics procedures show promise in treating immature teeth with necrotic pulp and apical periodontitis. This procedure involves the replacement of damaged and infected pulp tissue with viable tissue that restores the normal tooth structure and function. Antimicrobials are currently used to control the infection; however, they are cytotoxic to stem cells of the apical papilla (SCAP) and can lead to root canal calcification. Management of these teeth requires a scaffold that can control root canal infection, wick the blood into the canal, and support the viability and differentiation of SCAP while inhibiting intracanal calcification. This study aims to develop a composite scaffold made of polyphosphate, a calcium binding inorganic polymer shown to promote cell proliferation and tissue regeneration, chitosan, a natural antimicrobial polymer that supports stem cell viability and activity, and copper, a metal ion with bactericidal properties.

**Methodology:** The scaffold was prepared by adding copper (Cu) to chitosan solution, followed by polyphosphate. The resulting scaffold was then freeze-dried and analyzed for elemental composition, chemical structure, release of Cu, antibacterial properties, cytotoxicity, as well as differentiation and mineralization assays. Data were analysed by a two-way analysis of variance (ANOVA) followed by the Tukey post hoc test.

**Result:** This study demonstrates that, by combining polyphosphate and chitosan, we could fabricate a scaffold that inhibits bacterial growth by 40 % and supports the viability of fibroblast and SCAP. Adding copper to this scaffold further increased bacterial growth inhibition by up to 68% while preserving cell viability. The immunocytochemistry and Alizarin Red staining revealed that this scaffold also supports the odontogenic differentiation of these stem cells while inhibiting their mineralization potential. Furthermore, this scaffold can be fabricated as a 3D cone-shaped scaffold with a strong vertical wicking ability at a rate of 0.5 mm/s and excellent degradability, with 53 % of the scaffold degraded after 28 days.

**Conclusions:** This study shows that a copper-loaded chitosan-polyphosphate scaffold combines biocompatibility, antibacterial activity, wicking ability, and biodegradability and has great potential as an endodontic regenerative scaffold.

## 1. Introduction

Endodontic treatment of immature teeth with pulp necrosis and apical periodontitis is particularly challenging due to their open apices and thin, fragile dentinal walls [1], [2]. They are conventionally treated by apexification, in which either calcium hydroxide is repeatedly applied over several months to induce apical barrier formation or, more recently, Mineral Trioxide Aggregate plugs are utilized to provide an immediate apical barrier [3] [4]. However, apexification techniques do not promote additional root development, and the tooth remains non-vital and susceptible to fracture [3]. To overcome these limitations, regenerative endodontics (RE) has been introduced to promote pulp tissue regeneration and root maturation.

RE protocols involve minimal instrumentation, extensive but cautious irrigation, and placement of an intracanal medication for further disinfection. At the second appointment, intracanal bleeding is stimulated, followed by the formation of a blood clot inside the root canal [3], [5], [6]. This protocol aims to disinfect the root canal, resolve clinical symptoms, and recruit stem cells from the apical papillae (SCAP) directly next to the root tip to regenerate the dentine-pulp complex and promote further root maturation [7]. However, there is still uncertainty about the clinical outcome of regenerative endodontics (RE) due to persistent infection and calcification of root canals, which may eventually cause root canal obliteration that interferes with the normal functioning of dental pulp [8]. It has been reported that 79% of failed RE cases are caused by persistent infection [9], while intracanal calcifications occur in a frequency range of 20% to 91% [10] depending on whether the underlying cause is the medicament used or the bleeding induced during the procedure [8].

Usual intracanal medicaments for RE include calcium hydroxide and double or triple antibiotic pastes. Calcium hydroxide and double antibiotic pastes have less antimicrobial efficiency than triple antibiotic paste; however, a triple antibiotic paste is toxic to stem cells [9]. Any remaining bacteria may inhibit the differentiation and regeneration of SCAP [11]. Moreover, calcium hydroxide and antibiotic paste treatment increased intra-canal calcification incidence to 76.9% and 46.2%, respectively [10]. Research findings have indicated that the use of biocompatible and biodegradable scaffolds with inherent antimicrobial properties can aid in the success of regenerative endodontics [11], [12] .

Various biologically derived, naturally derived, and synthetic scaffolds have been studied for the delivery and growth of SCAP in the root canal space [13]. Among naturally sourced scaffolds, chitosan (Chi), a polysaccharide biopolymer derived from the N-deacetylation of chitin, is very promising due to its broad antimicrobial spectrum, biocompatibility, and biodegradability. [13 - 15]. Moreover, the chitosan structure is similar to components of the extracellular matrix (3), and a recent in vivo study did show its ability to promote the biological activities of dental pulp stem cells [15]. However, the antimicrobial action of chitosan scaffolds is limited to certain species of bacteria [17]. Several studies have been conducted to enhance the antimicrobial effectiveness of chitosan scaffolds by incorporating antimicrobial therapeutic ions [17–20].

Copper (Cu) is a powerful antimicrobial therapeutic ion that is effective against a broad spectrum of pathogens including *E. faecalis* [18], [21], [22], the most prevalent species in persistent endodontic infections that is also highly resistant to antimicrobial agents [23], [24]. Additionally, copper enhances vascularization and promotes tissue ingrowth within the scaffold [20]. Chitosan chains are known to chelate with copper ions[18] [19]; however, copper is released from the chitosan-copper complex within the first 10 hours after application [21]. While a high concentration of copper ions can have a strong antibacterial effect, it can also result in cytotoxicity [19]. Consequently, a sustained and slower release of copper ions at the appropriate antibacterial concentration is necessary.

Chitosan and copper can both be bound to polyphosphates (PP), an inorganic polyanion polymer that consists of phosphate groups linked by oxygen molecules [17], [25], [26]. PP is biocompatible, biodegradable, and has purported bactericidal properties, particularly against *P. gingivalis* associated with periodontal and endodontic infection [27], [28] . It is also known as a potent mineralization inhibitor, impeding ectopic calcification by direct calcium chelation rather than as a result of changes in cell or matrix [29]. Accordingly, we hypothesize that a copper-loaded chitosan-polyphosphate (Chi-PP) scaffold can support the viability and odontogenic differentiation of SCAP while preventing intracanal calcification and providing sustained and controlled release of copper for short-term infection control.

To test our hypothesis, we first optimized the processing conditions to produce a porous copper-loaded Chi-PP scaffold. The scaffold was then evaluated for its biocompatibility, antimicrobial effectiveness against *E. faecalis*, and clinical applicability with respect to its degradation rate, vertical wicking capacity, and ability to shape.

## 2. Materials and Methods

The manuscript of this laboratory study has been written according to Preferred Reporting Items for Laboratory studies in Endodontology (PRILE) 2021 guidelines (Supplementary Fiigure 1) [30].

### 2.1. Synthesis of Copper-Loaded Chi-PP Scaffold

Sodium polyphosphate with a degree of polymerization of 22 was prepared using baseline processing protocols previously established in the Filiaggi lab [31]. Medium molecular weight chitosan (Glentham Life Sciences) with a degree of deacetylation (DDA) of ≥ 90% and molecular weight of 125,000 kDa was used to prepare copper-loaded Chi-PP scaffolds. Firstly, chitosan powder was dispersed in 1% glacial acetic acid and stirred for 4 hours to achieve dissolution at 15 mg/mL, while PP solution was prepared in deionized water at a concentration of 15 mg/mL. Copper chloride (CuCl_2_.2H_2_O, Sigma-Aldrich) was added to the chitosan solution at three different concentrations to achieve approximate saturation of 3% (Cu1), 6 % (Cu2), and 12% (Cu3) of the free amino groups of chitosan[18] and left to disperse homogeneously for one hour.

Subsequently, the Chi-Cu solution was added to the PP solution at a volume ratio of 1 and the mixture was immediately mixed for 2 minutes at high-speed using a homogenizer (Fisher Scientific). The resulting mixture (polyelectrolyte complex) was then left to settle at 4 ° C overnight prior to rinsing with TRIS buffer (50 mM, pH 7.4) using a vacuum filter to neutralize the pH. The mixture was subsequently inserted in a conical plastic mold and frozen and freeze-dried overnight using a freeze dryer (Labconco® FreeZone® 2.5 Liter). The control Chitosan group was prepared without the addition of copper or PP. To evaluate the impact of PP on copper release, two groups of Chi-Cu (Chi-Cu1 and Chi-Cu3) were established to study copper release before and after the addition of PP. For ease of handling, the Chi, Chi-Cu1, and Chi-Cu3 groups were frozen and freeze-dried before and after washing. Overall, samples for seven experimental groups in total were fabricated (Table 1).

**Table 1:**
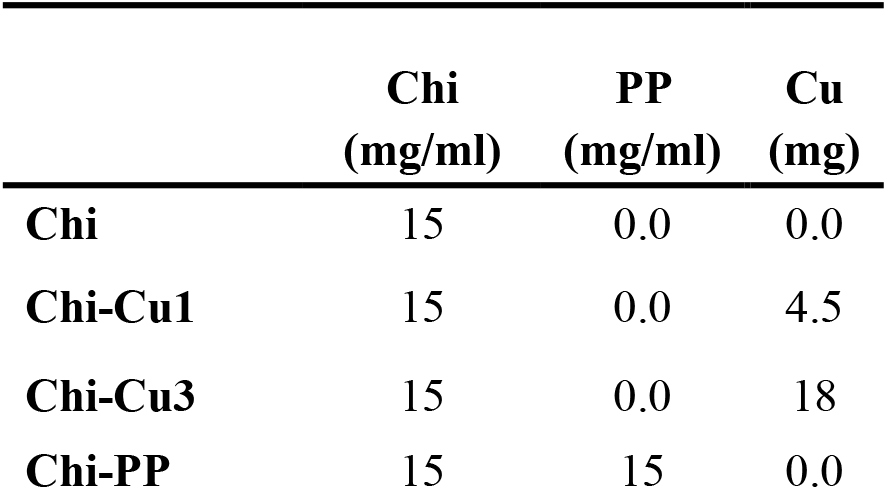
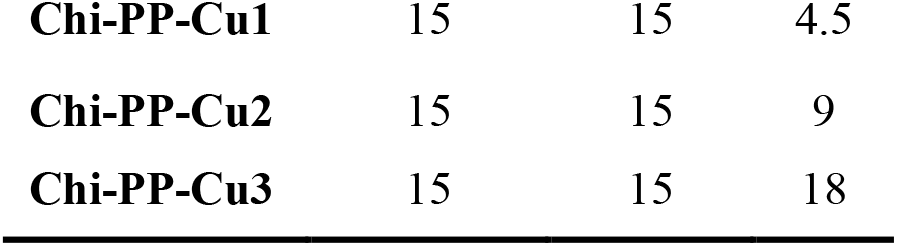
Concentration of initial reactants for fabricating different scaffolds and their corresponding labeling.

### 2.2. Scaffold Characterization

#### 2.2.1. Chemical and structural analysis

The concentration of phosphorus (P) and copper (Cu) in the resulting samples was determined with Inductively Coupled Plasma-Optical Emission Spectroscopy (ICP-OES; Perkin Elmer Optima 8000). Approximately 10 mg of samples (n=3) was accurately weighed, transferred to a 50 mL Falcon tube, and dissolved in concentrated nitric acid (6 mL) and HCL (3 mL) at room temperature. After complete sample digestion, a high-purity deionized water was added to adjust the final volume to 50 mL. 2 mL of the resulting solutions were further diluted into deionized-distilled water (8 ml) prior to submitting for ICP-OES analysis. Quality control checks were done prior to the initial analysis and every 20 consecutive samples.

The interaction between phosphoric and ammonium ions of PP and Chi, respectively, as well as the complexation of copper ions with both PP and Chi, were verified by Attenuated Total Reflectance Fourier Transform Infrared Spectrometry (ATR-FTIR; Bruker Vertex 70). The spectra were recorded in absorbance using a wavelength range of 400 – 4000 cm^−1^ at a resolution of 1 cm^−1^. The resulting spectra were analyzed using OPUS software.

#### 2.2.2. Copper release measurements

10 mg samples (n=3) were incubated in 5 mL of TRIS buffer (50 mM, pH 7.4) containing 16 µg/mL lysozyme (from chicken egg white, sigma) at 37°C. *In vivo*, chitosan is mainly degraded by lysozyme targeting the acetylated part [29]; therefore, lysozyme has been added to the immersion media in concentration like serum level (16 µg/mL). Aliquots were withdrawn from the media at regular intervals (3 hours and 1, 3, and 7 days), and the samples were re-immersed in fresh buffer at each collection time. The concentration of the released Cu from different scaffolds was then quantified using ICP-OES, as described in section 2.2.1.

#### 2.2.3. *In-vitro* degradation rates

10 mg samples were incubated in 5 mL of TRIS buffer (50 mM, pH 7.4) containing lysozyme (16 µg /mL) under stirring at 37 °C for 7, 14, 21, and 28 days, with media exchanges every other day. At each designated time point, the samples (n=3) were centrifuged to remove the media, dried at ambient temperature, and weighed to calculate the weight loss. Degradation rate was reported as the % of weight loss according to the following equation:

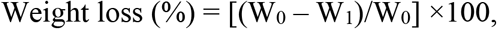

where W_0_ and W_1_ are the weights of the samples before and after incubation, respectively.

#### 2.2.4. Vertical wicking measurements

The wicking level and speed within different scaffolds (n=3) were determined using Dulbecco’s Modified Eagle Medium (DMEM) containing 0.02% phenol red for observing the height of the media wicked into the scaffold. All wicking measurements were carried out at 37°C. The scaffolds (with a length of 30 mm) were inserted into a plastic cone-shaped mold with an open tip. The tip end of the mold was then immersed in 0.2 mL of DMEM and kept vertically beside a ruler, as shown in the Supplementary Fiigure 2. The time required for media to pass through each scaffold was recorded until a wicking distance of 20 mm was achieved. The vertical wicking rate was reported as mm/s.

### 2.3. Scaffold Cytotoxicity

The cytotoxic concentration of copper-loaded scaffolds was determined using a 3T3-L1 cell line. Different scaffold formulations (40 mg each, n=3) were sterilized for 1 hour in a UV oven to confirm microbiological sterility. DMEM with 10% fetal bovine serum and 1% antibiotic-antimycotic solution was conditioned with the scaffold formulations at 40 mg/mL concentrations for 24 hours in an orbital incubator at 37°C. Subsequently, conditioned media was extracted by centrifuging and diluted by 2 fold. Cells were seeded on a 48-well plate at 40% confluency (8×10^3^ cells/well) and incubated overnight at 37°C and 5% CO_2_ (Three independent cultures with three samples each.), followed by exposure to various scaffold extracts (200 µL/well) for an additional 24 hr. After treatment, 3T3-L1 viability was evaluated using an Alamar Blue assay and a Live/Dead assay, which were performed according to the manufacturer’s instructions; for Live/Dead assays, cells were visualized using an EVOS FL Auto 2 microscope (Invitrogen, Waltham, MA). Cell viability of scaffold treatment groups was normalized against the scaffold-free DMEM (blank control).

### 2.4. Odontoblastic Differentiation and Mineralization Potential

These experiments utilized SCAP expressing STRO-1 isolated from immature third molars, which had previously been purified through immunomagnetic separation as described by Raddall et al [3]. The use of SCAP was approved by the Nova Scotia Health Research Ethics Board (REB FILE #: 1022150). SCAPs were seeded in α-minimum essential medium (MEM) supplemented with 10% fetal bovine serum, 1% L-glutamine, and 1% antibiotic-antimycotic solution using a 6-well plate at 10% confluency (25×10^3^ cells/well) and incubated overnight at 37°C and 5% CO_2_. To induce odontogenic differentiation, the scaffold (Chi, Chi-PP, or Chi-PP-Cu1) conditioned MEM media were supplemented with 0.01 mM dexamethasone disodium phosphate, 1.8 mM monopotassium phosphate, and 5 mM β-glycerophosphate. Positive and negative control samples consisted of SCAPs grown in scaffold-free MEM medium for the same period with or without any odontoblast differentiation factors, respectively. SCAP cells were exposed to scaffold-conditioned or scaffold-free media (2 mL/well) for 7, 14, and 21 days at 37°C and 5% CO_2_, with a media change every second day. SCAP viability was evaluated after 7 days using Alamar Blue and Live/Dead assays as described above.

#### 2.4.1. Alizarin red staining for mineralized matrix

Extracellular calcium deposits from differentiated odontoblasts were detected after 7, 14, and 21 days using Alizarin Red S staining (Sigma Aldrich). Alizarin Red S powder was dissolved in distilled water at a pH of 4.2. Cells were washed with PBS, fixed with 4% paraformaldehyde (PFA) for 30 min, and washed with distilled water. Subsequently, Alizarin Red S staining solution was added to each well for 1 hr at room temperature in total darkness. Thereafter, wells were washed with deionized water and PBS was added. The cells were viewed under a bright-field EVOS FL Auto 2 microscope (Invitrogen, Waltham, MA).

#### 2.4.2. Immunocytochemistry to assess odontogenic differentiation potential

The odontogenic differentiation potential of SCAPs was evaluated after 7 days using immunocytochemistry staining of dentin matrix protein-1 (DMP-1) and dentin sialophosphoprotein (DSPP). These proteins are specific markers of odontogenic differentiation expressed in odontoblasts but not in undifferentiated SCAPs [3]. Cells were washed with PBS and fixed with 4% paraformaldehyde at 4°C overnight. To permeabilize the cells, cells were rinsed with PBS and incubated in 0.2% (w/v) Tritan-X100 at room temperature for 4 min. Subsequently, the fixed cells were incubated with either rabbit anti-DMP-1 or rabbit anti-DSPP (Abcam, Cambridge, MA) primary antibodies diluted 1:250 in incubation buffer overnight at 4°C. Secondary donkey anti-rabbit immunoglobulin G Alexa Fluor 647 (Abcam) was diluted 1:1000 in 1% bovine serum albumin and incubated with all samples for 2 hours at room temperature, followed by 40,6-diamidino-2-phenylindole (DAPI) staining solution (Abcam) diluted 1:1000 in PBS and Phalloidin-iFluor 488 reagent (Abcam) diluted 1:1000 in PBS for 20 minutes. The labelled cells were visualized and imaged under an EVOS FL Auto 2 fluorescent microscope (Invitrogen, Waltham, MA).

### 2.5. Determining the Antimicrobial Effectiveness of the Scaffold

The antimicrobial properties of the scaffold against *E. faecalis* were evaluated using a colony-forming unit assay. Briefly, *E. faecalis* were streaked on agar plates consisting of 25 ml LB broth and 1.5% w/v of agar and inoculated at 37°C for 24 hr. One colony of *E. faecalis* was added to 5 mL of LB media and inoculated at 37°C for 24 hr. *E. faecalis* cultures were adjusted to an optical density of 0.1 and added to the scaffolds (40 mg each) at a v/w ratio of 1/20 and inoculated at 37°C for 1, 3, 7, and 14 days; scaffold-free cultured media was used as a control. Every 24 hours, 1 mL of the cultured media was collected and replaced with LB fresh media, and the collected samples were re-incubated at 37°C. After serial dilutions of the collected media, 100 μL of each dilution were plated on an agar plate for *E. faecalis* and incubated overnight at 37 °C. Colonies were counted and calculated as percentages of remaining viable microorganisms relative to the negative control (scaffold-free control). The sample size was estimated based on studies comparing different scaffold formulations.

### 2.6. Statistical Analysis

Statistical analysis of Cu and P content, cell viability, bacterial growth inhibition and degradation and wicking rate was determined by a two-way analysis of variance (ANOVA) followed by the Tukey post hoc test using origin (OriginPro 9.0, Northampton, USA). Differences were considered significant when P < 0.05.

## 3. Result

### 3.1. Synthesis of a Copper-Loaded Chi-PP Scaffold with Controlled Copper Release

In this work, highly porous copper-loaded Chi-PP scaffolds were obtained through polyelectrolyte complexation of polyphosphate and chitosan containing different concentrations of Cu, followed by lyophilization.

The P and Cu content as determined by ICP-OES for complexed Chi-PP with varying Cu loading is shown in Figures 1a and b. A predictable increase in Cu loading in the scaffold correlating with increased Cu loading in the Chi solution was observed (Figure 1a). After scaffold washing and lyophilization, only 0.066±0.025, 0.13±0.003, and 0.26±0.002 mg of the loaded copper remained within Chi-PP-Cu1, Chi-PP-Cu2, and Chi-PP-Cu3 scaffolds, respectively. However, significantly (p<0.0001) more copper was incorporated within scaffolds prepared without PP, where 0.13±0.006 and 0.45±0.01 mg of the loaded copper were incorporated within Chi-Cu1 and Chi-Cu3, respectively. On the other hand, a significantly (P<0.001) higher proportion of phosphate ions was incorporated into Chi-PP-Cu1 (0.96 ±0.08 mg), Chi-PP-Cu2 (1.045 ±0.02 mg), and Chi-PP-Cu3 (1.045±0.006 mg) compared to Chi-PP (0.77±0.07mg), with no significant difference between Chi-PP-Cu1, Chi-PP-Cu2, and Chi-PP-Cu3, P>0.17. (Figure 1b).

**Figure 1:**
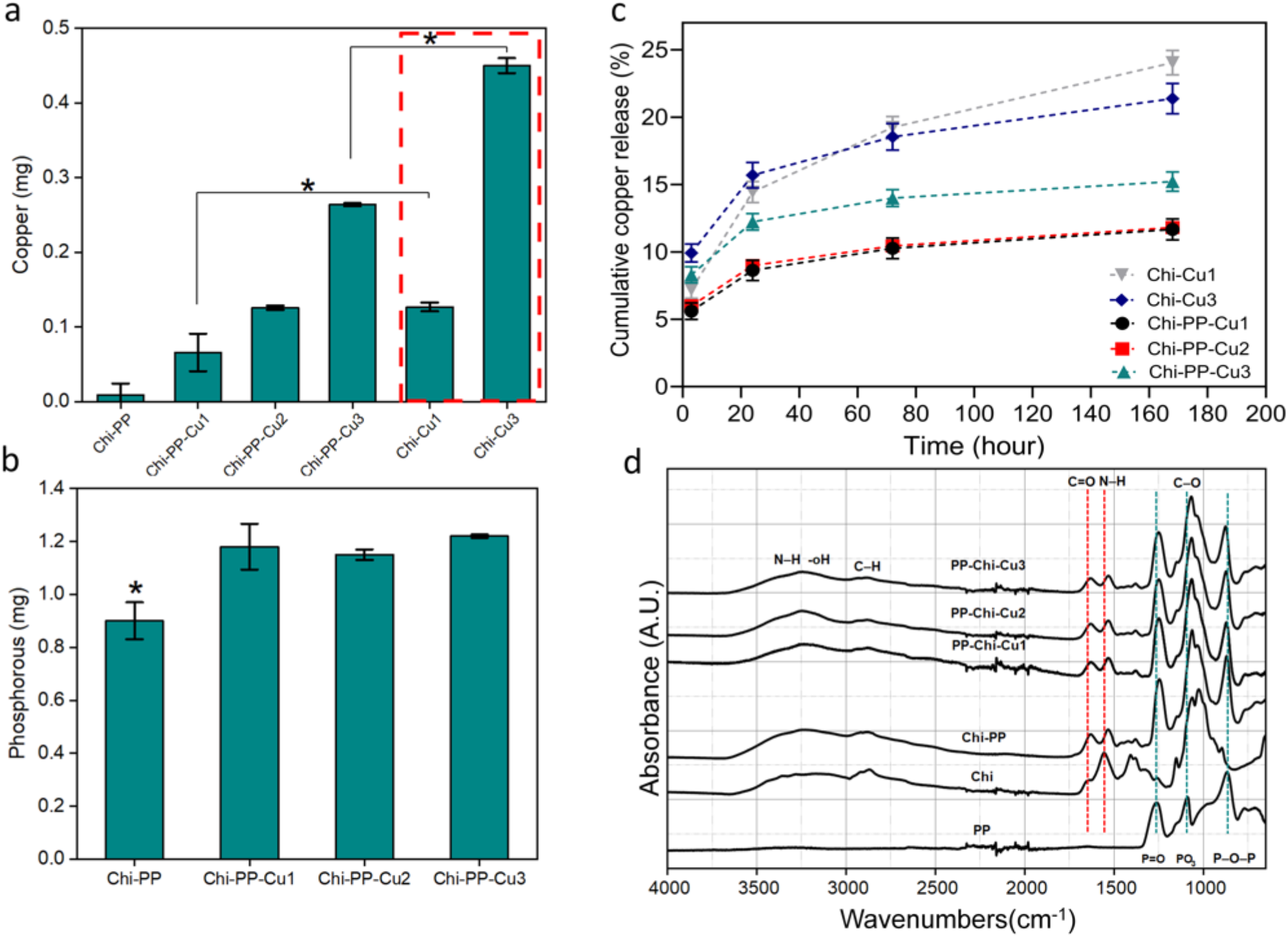
(a) Cu and (b) P content as determined by ICP-OES for complexed Chi-PP with varying Cu loading. A higher proportion of phosphate ions was incorporated into scaffolds containing copper, * P<0.001 indicates a significantly lower phosphate content in copper-free scaffold, and a predictable “dose-dependent” response (increased Cu loading in the scaffold correlating with increased Cu-loading in the Chi solution) was observed, * P<0.0001 indicates a significantly more copper content in PP-free scaffold. (c) Cumulative copper release from all copper-loaded scaffolds as a function of time. (d) ATR-FTIR spectra of Chi, PP and complexed Chi-PP with varying Cu loading.

The results from Cu release studies are shown in Figure 1c. After 24 hours, around 9 % of the loaded copper was released from the Chi-PP-Cu1 and Chi-PP-Cu2 samples, and 12 % was released from the Chi-PP-Cu3 samples. During the next 7 days, an even slower release of copper was observed in these groups, with only 3% more copper being released. On the other hand, samples prepared without PP (Chi-Cu1 and Chi-Cu3) released about 15% of their copper content within 24 hr and an additional 10% from Chi-Cu1 and 6% from Chi-Cu3 over the next 7 days.

### 3.2. Confirmation of Complexation by ATR-FTIR

ATR-FTIR spectra of the Chi-PP scaffold before and after copper loading are represented in Figure 1d along with the spectra of each precursor polymer (chitosan and PP) used as controls. The absorption spectrum of PP exhibited the following characteristic bands: stretching of the P-O-P bridge at 863 cm^-1^; stretching of P = O at 1262 cm^-1^; stretching vibrations in PO_3_ at 1088 cm^-1^; and stretching of the P-O-P bridge at 863 cm^-1^[32]. After chitosan addition, the positions of these bands that characterized the middle of the polyphosphate chain (P-O-P; P=O) and the terminal atoms (PO_3_) were displaced to 868, 1242, and 1062 cm^-1^, respectively. This result is consistent with previous observations on a chitosan-polyphosphate system, suggesting the potential for chitosan chain interactions with PP at internal phosphate units as well as at terminal sites. [33], [34].

The chitosan spectrum exhibited characteristic absorption bands at 1652 cm^-1^ and 1551 cm^-1^, corresponding to C=O stretching and NH_2_ bending vibration in the amide group[32]. The 1551 cm^-1^ peak is sharper than the peak at 1652 cm^-1^, which indicates the high degree of deacetylation of the chitosan. After complexation with PP, the intensity of the (C=O) band at 1652 cm^-1^ decreased dramatically, while the intensity of the (NH_2_) band at 1551 cm^-1^ increased. A significant displacement to a lower wavenumber was also noted, with the band at 1652 cm^-1^ shifted to 1629 cm^-1,^ and the band at 1551 cm^-1^ shifted to 1526 cm^-1^. These changes suggest that the ammonium groups of chitosan interacted with polyphosphate, as previously reported by D. R. Bhumkar and V. B. Pokharkar1 [35].

Copper loading induced further changes in the characteristic FTIR spectra and bands of the Chi-PP scaffold. In particular, a broad band in the region of 3500–3200 cm^−1^ corresponding to the stretching vibration of NH2 and OH that are involved in hydrogen bonds [34] was noticeably less broad when copper was loaded. In addition, there was a significant decrease in the intensity of the C–O stretching band at 1024 cm^−1^, which is present in pure chitosan. These modifications to the copper-loaded Chi-PP spectrum may suggest a possible interaction between amino and carbonyl groups of chitosan and copper, as reported in a previous copper-chitosan study [36]. The spectrum of copper-loaded Chi-PP scaffolds also exhibited a slight displacement of the P-O-P stretching bands at 868 cm^-1^ in Chi-PP spectra to 871 cm^-1^ and 875 cm^-1^ in Chi-PP-Cu2 and Chi-PP-Cu3, respectively. The band located at 1242 cm^-1^, which is associated with the (P=O) band of the PP chain in Chi-PP, was found to have shifted to 1248 cm^-1^ in copper-loaded groups. This displacement suggests that copper ions and /or copper chelating with Chi are bound to the PP chain.

### 3.3. The Cytotoxic Effect of the Scaffolds on 3T3-L1 and SCAP Cells

Based on the literature, copper has a minimum inhibitory concentration (MIC) of 10 ppm, while its *in vitro* cytotoxic concentration is 30 ppm [21]. The scaffold is expected to exhibit antibacterial activity without any cytotoxic effects within the range of 10-30 ppm. In order to verify this, the concentration of copper released from the copper-loaded Chi-PP scaffolds was compared to the literature-reported values and presented in Figure 2a. Based on the results, it was confirmed that adjusting the amount of copper initially loaded in the chitosan can precisely control the release. Specifically, the Chi-PP-Cu1 scaffold released about 10.5 ± 1.15 ppm of copper within the first 24 hours. This concentration is within the optimal range that can prevent bacterial colonization without causing cytotoxicity, in contrast to Chi-PP-Cu2 and Chi-PP-Cu3, which released 22.63 ± 1.04 and 51.22 ± 8.1 ppm of copper after 24 hours, respectively.

**Figure 2:**
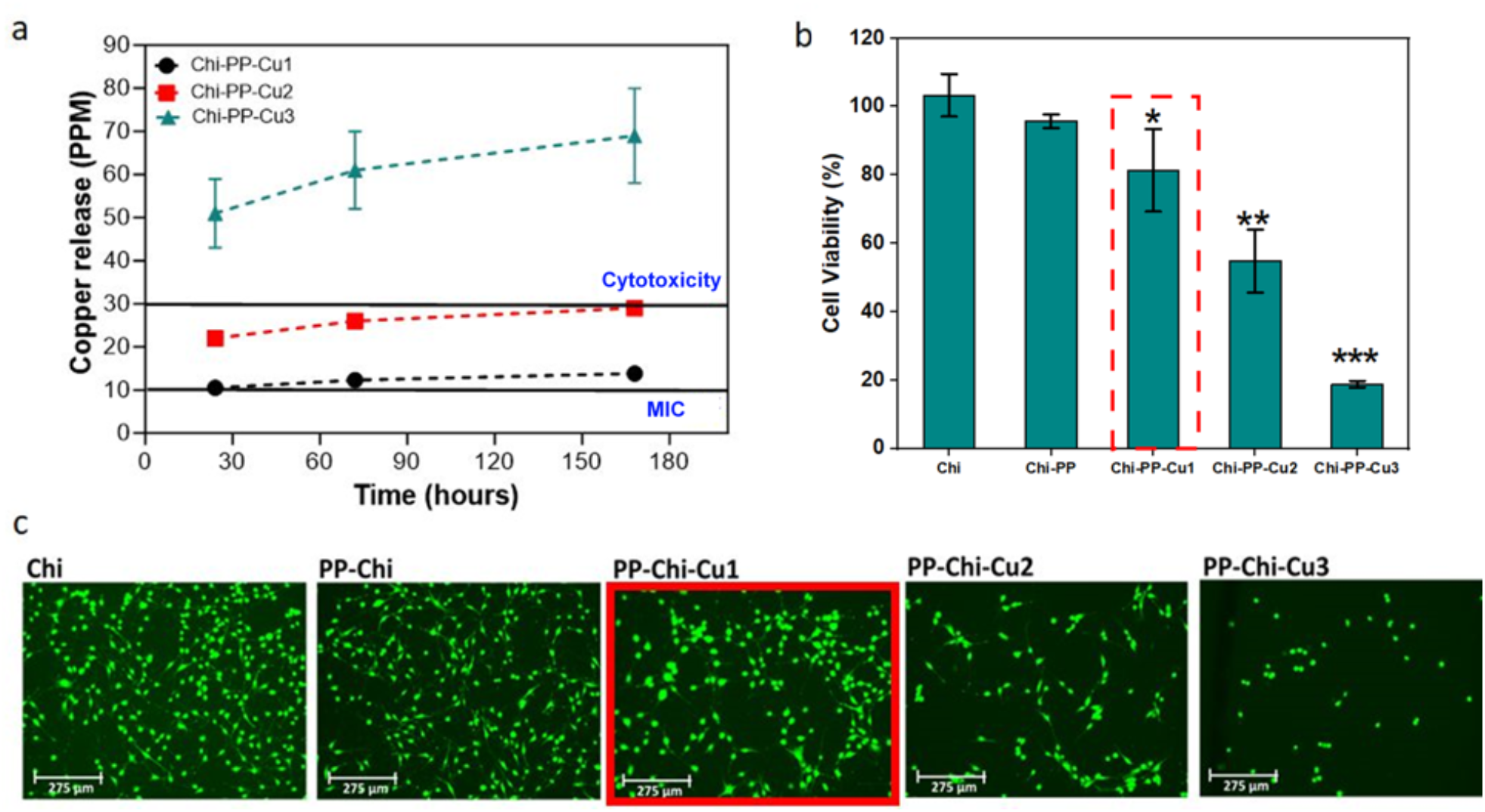
(a) Concentration of released copper over time, the minimal inhibitory concentration (MIC, 10 ppm), and cytotoxic concentration (30 ppm) reported in the literature for copper are plotted side by side for comparison. (b) Effect of Cu on L1-3T3 cell viability after 24 h of exposure to different scaffold formulations using Alamar blue assay, Chi-PP-Cu1 scaffold (highlighted with the red dotted line) presented the highest Cu concentration that preserved cell viability; (c) L1-3T3 cell viability (Live/Dead assay), representative images of L1-3T3 cell lines stained with calcein AM (live, green) and ethidum homodimer (dead, red) viability staining technique after a 24 h of treatment with different scaffold formulations or without treatment (positive control). The scale bar at the left lower corner is 275 µm. Asterisk (*) indicates significantly lower cell viability as copper concentration increased, * P<0.04, ** P<0.0002, *** P<0.0001.

Subsequently, an Alamar Blue assay was used to evaluate the viability of 3T3-L1 cells in the presence of different scaffold extracts. The results were compared to the blank control and expressed as a percentage difference, as shown in Figure 2b. After 24 hours, Chi and Chi-PP extracts were found to be non-toxic with 103.2 ± 6.2% and 95.7 ±1.9 % cell viability, respectively. However, when loaded with copper, the Chi-PP scaffolds showed a significant dose-dependent decrease in cell viability. The viability of cells was 81.3 ±12 %, 54.8 ± 9.2%, and 18.7± 0.95 % with Chi-PP-Cu1, Chi-PP-Cu2, and Chi-PP-Cu3, respectively. According to ISO 10993-5, an extract is considered cytotoxic if its cell viability is reduced to more than 30% of the blank control [37]. Although Chi-PP-Cu2 released just above 20 ppm of copper within the first 24 hours, it was highly toxic to cells. As a result, only the non-cytotoxic Chi-PP-Cu1 scaffold was used for further experiments. The viability of 3T3-L1 cells was also tested using the Live/Dead assay, which confirmed the Alamar Blue assay results (Figure 2c).

The viability of SCAP was evaluated using Alamar blue and Live/Dead assays after being subjected to extracts from the Chi-PP-Cu1 scaffold as well as a Cu-free (Chi-PP) and Chi-only (Chi) scaffold in odontogenic differentiation media for 7 days. Alamar blue assay revealed that both Chi and Chi-PP extracts showed no toxic effects, with 94.8 ± 13.8 % and 106 ± 0.86 % cell viability, respectively. However, the Chi-PP-Cu1 extract exhibited a 72 ±3.8 % cell viability but was still non-cytotoxic as per ISO 10993-5 standard (Figure 3b). The Live/Dead assay results were consistent with those obtained from the Alamar Blue assay (Fig. 3a) regarding SCAP cell viability. Therefore, the study proceeded to evaluate the effects of copper-loaded Chi-PP scaffolds on SCAP cell differentiation and mineralization potential.

**Figure 3:**
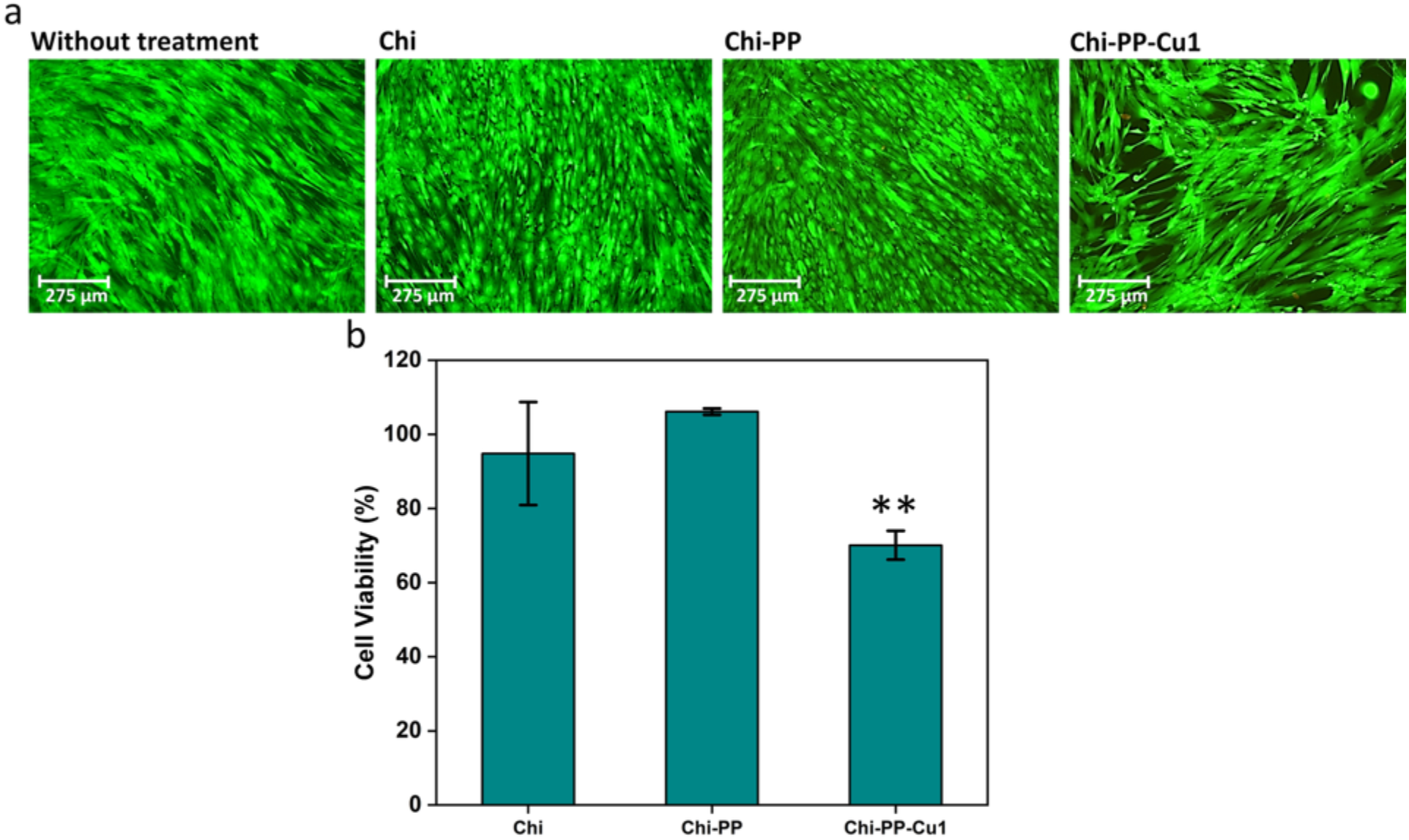
(a) SCAP cell viability (Live/Dead assay), representative images of SCAP cell lines stained with calcein AM (live, green) and ethidium homodimer (dead, red) viability staining techniques after 7 days of treatment with different scaffold formulations or without treatment (positive control). (b) Effect of different scaffold formulations on SCAP cell viability after 24 h of exposure to different scaffold formulations using Alamar blue assay. The scale bar at the left lower corner is 275 µm. Asterisk (*) indicates significantly lower cell viability compared to copper free groups. **P<0.00027

### 3.4. Odontoblastic Differentiation and Mineralization Potential of SCAP

In order to confirm the differentiation of SCAPs into odontoblasts, it is necessary to use an immunocytochemical staining technique to investigate the expression of specific markers such as dentin matrix protein-1 (DMP-1) and dentin sialophosphoprotein (DSPP). These markers are known to be specific for the odontogenic differentiation of SCAPs[38]. The expression of DMP-1 on STRO-1 SCAP cultured on odontogenic media without or with scaffold extracts, including Chi, Chi-PP, or Chi-PP-Cu1, was minimal and showed no significant difference between the groups (Fig. 5a-d). However, DSPP was highly expressed and showed a similar pattern across all groups (Fig. 5e-l).

**Figure 4:**
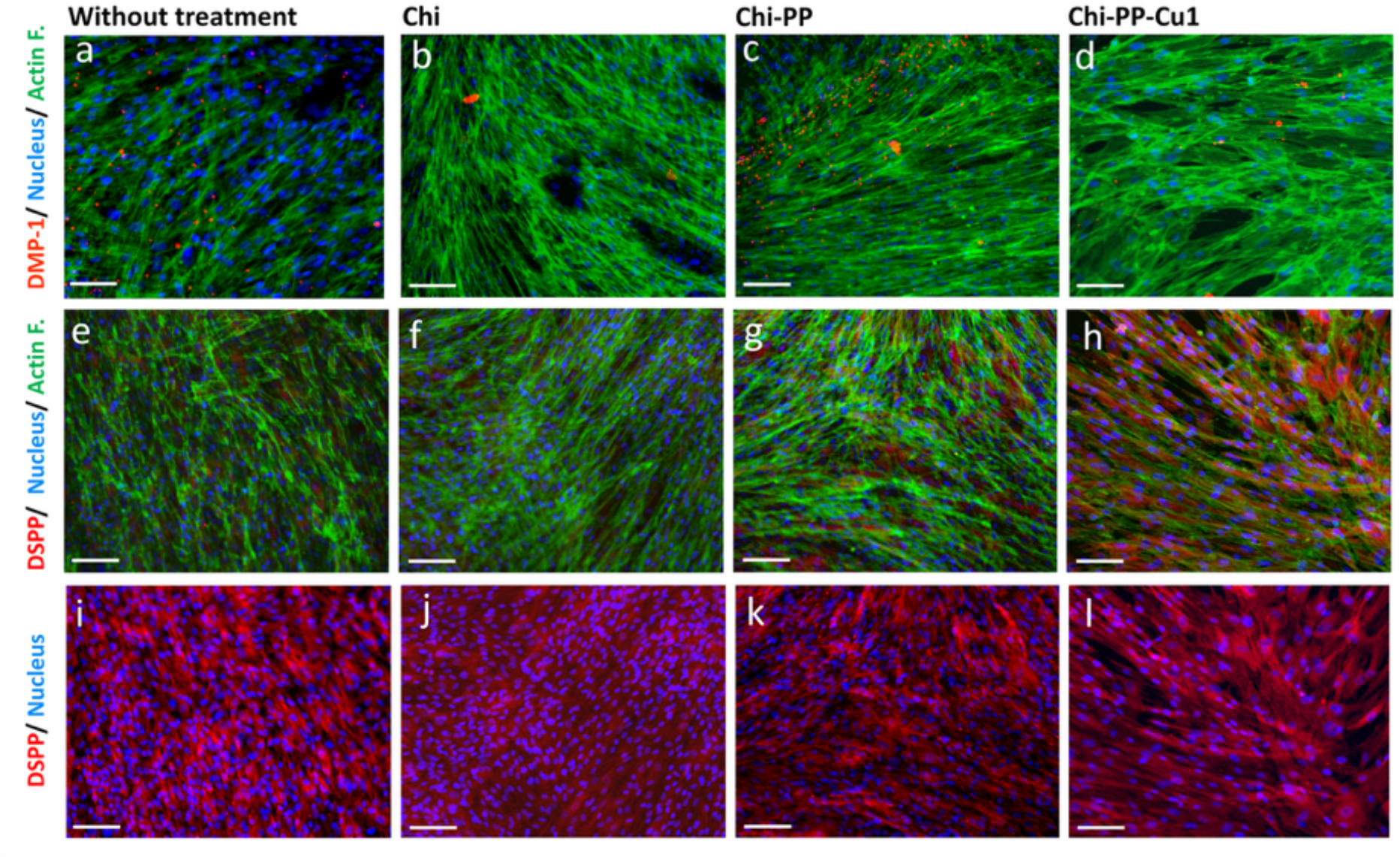
Immunocytochemical detection of the odontogenic differentiation markers (DMP-1 and DSPP). Representative images of SCAPs with immunocytochemistry staining for (a-d) DMP-1 protein (red stain) and (e-l) DSPP protein (red stain) after 7-day exposure to odontogenic medium without (positive control) or with exposure to different scaffold formulations. Nuclei were stained with DAPI, shown in blue, while F-actin was stained with Phalloidin, shown in green. The scale bar at the left lower corner is 100 µm.

**Figure 5:**
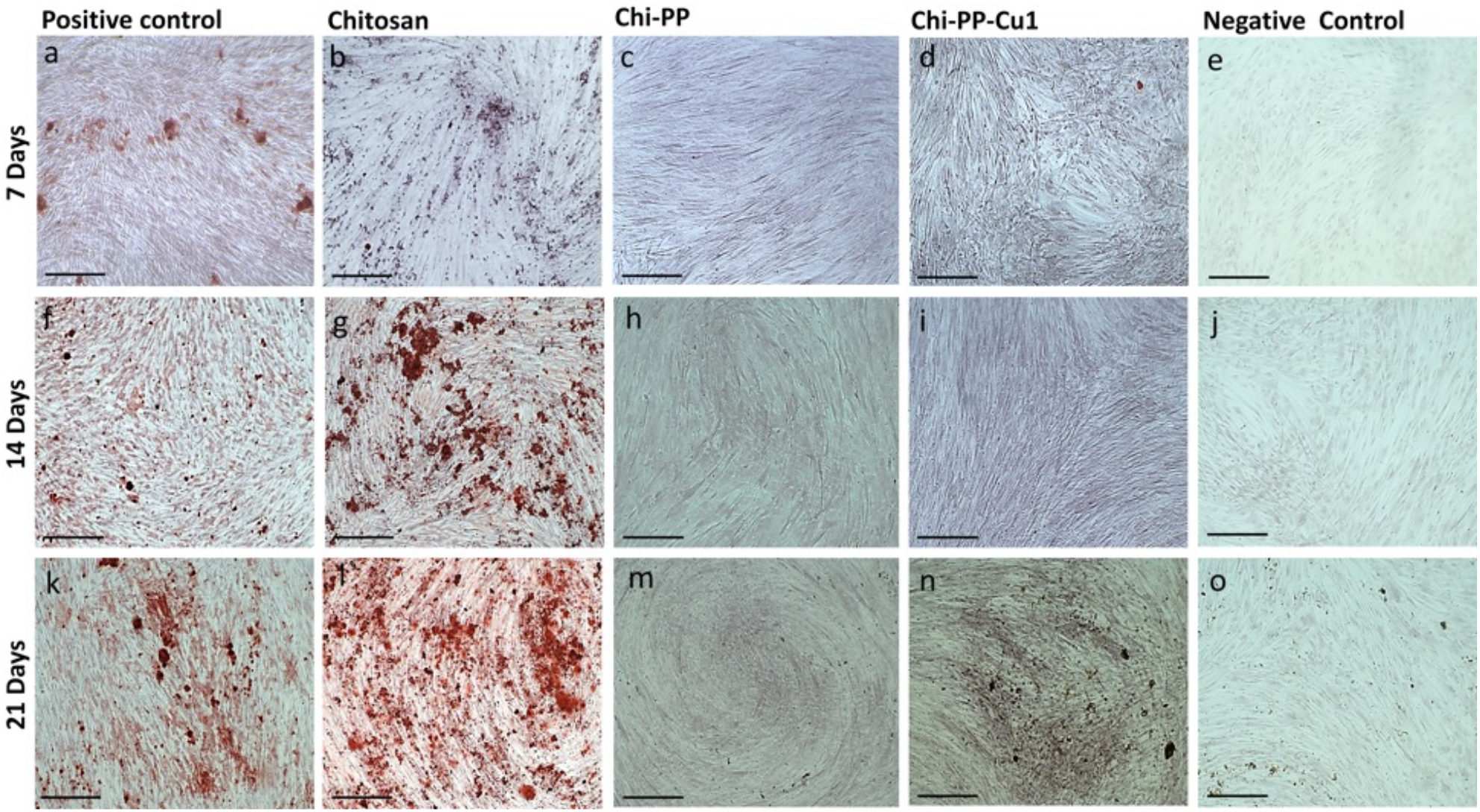
Microscopic images of Alizarin Red S staining of stem cells of dental papilla (SCAPs) cultured for 7 days (a-e), 14 days (f-g), or 21 days (k-o) in pure odontogenic medium (positive control), in odontogenic media treated with different scaffold formulations (Chi, Chi-PP, and Chi-PP-Cu1) or in regular α-MEM medium (Negative control). Calcium deposits (red color) are evident in SCAPs cultured in the presence of chi scaffold (b, g and l), but not in the presence of Chi-PP (c, h, and m) or Chi-PP-Cu1(d, I and n). The scale bar at the left lower corner is 500 µm.

Extracellular calcium deposits were detected using Alizarin Red staining. These deposits appear as a bright orange-red colour (as shown in Figure 5). The staining revealed that SCAPs cultured in an odontogenic medium could form mineralized red nodules after 7, 14, and 21 days (Fig. 5a, f, and k). Furthermore, when these cells were cultured in an odontogenic medium that contained Chi extract, mineralized nodules were also observed after 14 and 21 days (as shown in Fig. 5 g and l). However, in cultures containing PP extracts (Chi-PP and Chi-PP-Cu1), no bright red deposits were visible (as shown in Fig. 5).

### 3.5. Antibacterial Effectiveness Against *E. faecalis*

The results of colony-forming assays conducted at 24 hours showed that Chi has an antibacterial effect on *E. faecalis*, reducing bacterial growth by 39.8 ± 16% (Fig. 6a). While the addition of PP did not have a significant effect on bacterial growth, adding copper to the Chi-PP scaffold increased the inhibition of bacterial growth significantly. Increasing the copper concentration led to an even greater antibacterial effect, with a 59.3 ± 2.7, 73.2 ± 3.3, and 99.9 ± 0.5% reduction of bacterial growth observed after 24 hours in a Chi-PP-Cu1, Chi-PP-Cu2, and Chi-PP-Cu3 scaffolds, respectively. Since only Chi-PP-Cu1 scaffolds can maintain cell viability, the antimicrobial experiment was extended to 14 days in the presence of Chi-PP-Cu1, and bacteria colonies were counted at 3, 7, and 14 days. All scaffold formulations were able to decrease bacterial growth, but complete inhibition was not achieved. Chi-PP-Cu1 was found to be the most effective in reducing bacterial growth after 14 days, with a reduction rate of 68.5 ± 7.3 % (Figure 6b).

**Figure 4:**
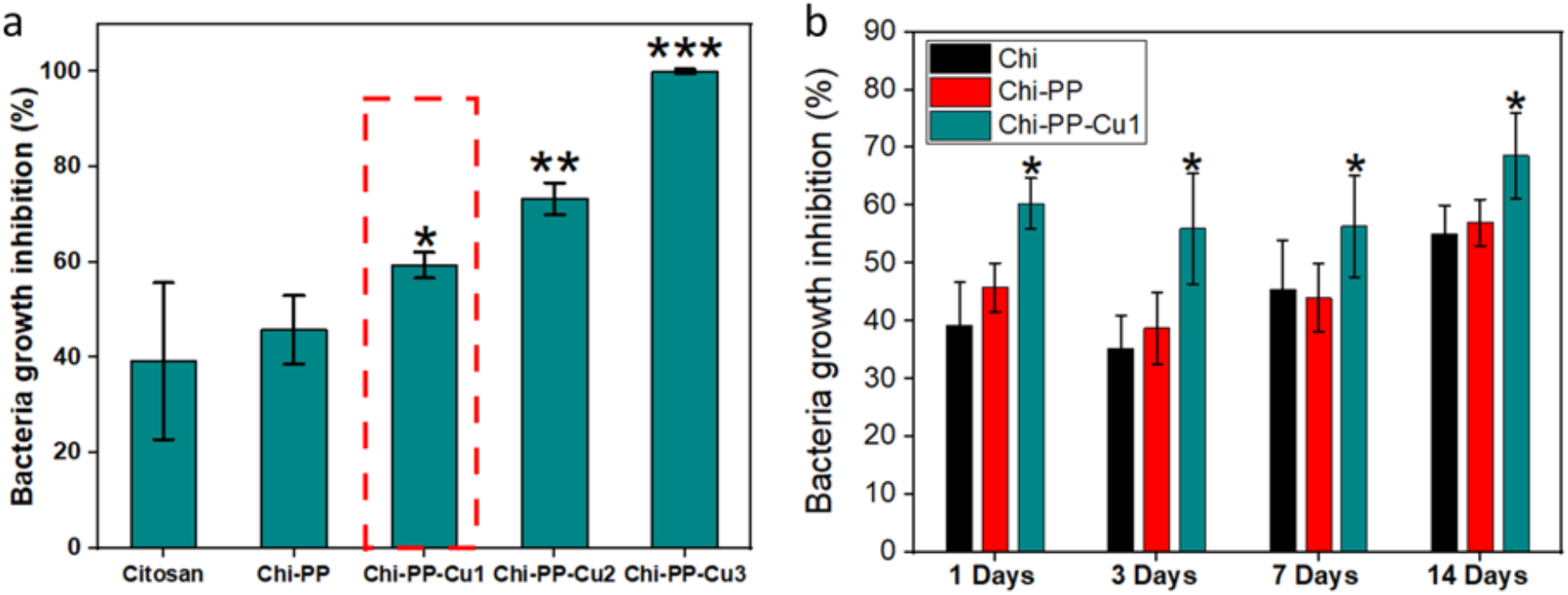
(a) Growth inhibition of bacteria (*E. Faecalis*) in a liquid medium after incubation with different scaffolds at 20 mg/ml concentrations for 24 h. Asterisk (*) indicates significantly higher bacterial growth inhibition as copper concentration increased, * P<0.0003, ** P<0.0002, *** P<0.0001.(B) Bacterial growth inhibition in a liquid medium after incubation with Chi-PP-cu1 at 20 mg/ml concentrations for 1, 3, 7, and 14 days. Asterisks (*) indicate a significantly (P < 0.0001) higher bacterial growth inhibition of copper-containing scaffold than that of copper-free groups.

### 3.6. The Clinical Applicability

A 3D conical shape, mimicking the dimensions and shape of dental roots, was intended for this scaffold in order to ease clinical application while supporting cell migration from the surrounding tissues. In this study, a qualitatively reproducible 3D conical scaffold was developed as shown in Figure 7a. Subsequently, the vertical wicking of liquid through the 3D conical scaffold was characterized to identify the rate and the height of liquid wicking along the scaffold. The results demonstrate that adding PP increased the rate and height of vertical wicking of media into the chitosan scaffold. The wicking rate of Chi-PP and Chi-PP-Cu1 was 0.5± 0.05 mm/s, considerably higher than the wicking rate of pure chitosan of 0.06 ±0.01 (Fig. 7b). Chi-PP and Chi-PP-Cu1 scaffolds were also able to wick fluid throughout their entire length (20 mm), while pure chitosan wicked only 5-8 mm of liquid.

The mass loss quantification of Chi, Chi-PP, and Chi-PP-Cu1 after soaking in lysozyme/Tris solution for 28 days is shown in Fig. 7c. The study found that pure chitosan scaffolds ‘experienced the least weight loss (18.3 ± 1.3 wt%) compared to other scaffolds. When PP and copper were added to chitosan, the degradation rate of the scaffold increased, with Chi-PP and Chi-PP-Cu1 exhibiting weight losses of 24 ± 1.2 wt% and 53 ± 5.3 wt%, respectively.

**Figure 5:**
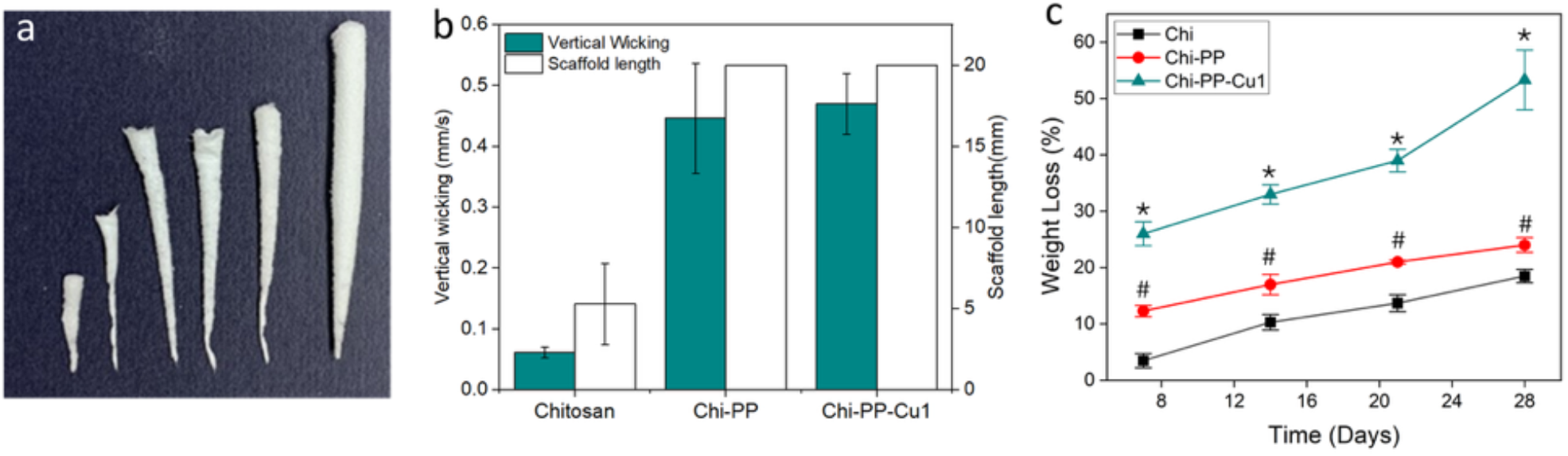
(a) Photograph of 3D cone-shaped copper-loaded Chi-PP scaffold achieved through polyelectrolyte complexation and lyophilization., (b) Vertical wicking rate and height, and (c) degradation rate of different scaffold formulations. Asterisks (*) indicate a significantly (P < 0.0001) faster degradation rate compared to the other groups and (#) indicate a significantly (P < 0.01) faster degradation rate than the Chi group and a slower degradation rate compared to the Chi-PP-Cu1 group.

## 4. Discussion

We present in this study the design and characterization of an innovative copper-loaded chitosan-polyphosphate scaffold in a formulation that promotes odontogenic differentiation of SCAP, while providing short-term infection control, inhibiting calcification, and degrading over time.

We first prepared chitosan and chitosan-copper solutions with three different concentrations of copper, and then we added polyphosphate and examined its effect on copper content and release. The structural analysis using ATR-FTIR confirmed that the polyelectrolyte (Chi and PP) complexation occurred and that copper ions and/or copper chelating with chitosan are bound to the polyphosphate chain. Therefore, the chitosan-polyphosphate scaffold loaded with copper is a hybrid composition of both organic and inorganic materials.

Elemental analysis of various formulations indicates that PP reduces both the amount of copper incorporated within chitosan scaffolds and its release, while scaffolds containing copper incorporate a higher proportion of phosphate ions. Based on these findings, it can be inferred that PP has the ability to regulate both the loading and release of copper. Copper seems to be interconnected between Chi and PP, which increases their interaction and decreased copper release rate. This implies that using PP with varying degrees of polymerization and concentrations could affect the loading and release of copper, which in turn may affect the antibacterial properties and cytotoxicity. Therefore, further research will be needed to explore the effect of different PP with varying degrees of polymerization.

Tissue engineering studies have revealed that both chitosan and polyphosphate are highly biocompatible [13, 27]. Cytotoxicity may occur, however, if copper ion concentrations exceed the optimal level. Cell culture results indicated that the Chi-PP composite was biocompatible and could support cell viability and proliferation. However, when loaded with copper, the Chi-PP scaffolds exhibited a significant, dose-dependent decrease in cell viability, with only the Chi-PP-Cu1 scaffold showing non-cytotoxicity. This finding is consistent with a previous Chi-Cu study that found that cell viability and copper concentration had a negative correlation, suggesting that copper ions are responsible for the decrease in cell viability [18]. This study has also shown that copper loaded Chi-PP scaffold supports odontogenic differentiation of SCAP. The expression analysis of DMP-1 and DSPP revealed that DMP-1 showed minimal expression while DSPP showed high expression. Based on the literature, it has been observed that DMP1 and DSPP genes show co-expression during the early stages of odontoblast differentiation before the initiation of mineralization. However, their expression patterns become significantly different during the later stages of differentiation. In particular, the expression of DMP-1 decreases in odontoblasts after the appearance of the minerals, while high levels of DSPP are sustained in odontoblasts [39], [40]. Hence, these findings suggest that the cells are well-differentiated and in the late stages of differentiation. However, it must be noted that cell culture studies cannot accurately replicate the conditions present in the human body. Therefore, it will be necessary to conduct *in vivo* studies using animal models to verify the scaffold’s biocompatibility and its effectiveness in tissue regeneration.

One of the limitations of regenerative endodontics is the calcification of root canals, which can lead to the obliteration of the root canal space, ultimately affecting the normal function of the dental pulp. Alizarin red staining was performed to qualitatively evaluates the deposited calcium on the presence of different fabricated scaffolds. Our findings showed that adding PP to chitosan prevent calcium deposits formation. This result is consistent with previous studies that have demonstrated that PP can impede mineralization in cell-free [41] and osteoblast cell culture models [29], as well as in induced aortic calcification model [42]. These studies have suggested that this inhibition occurs because PP binds with available calcium and inhibits tissue non-specific alkaline phosphatase without affecting cell differentiation or matrix secretion. This is consistent with the findings of this study, which found that PP did not affect odontogenic differentiation or cell viability, indicating that it inhibits mineralization via calcium chelation rather than via cell effects.

The success of regenerative endodontic therapy depends on the complete removal of microorganisms from infected root canals [9]. Chitosan and copper, known for their antibacterial properties, could be effective in inhibiting remaining bacteria that may hinder the regenerative process. Various types of chitosan solutions, gels, and nanoparticles have been studied in order to determine whether they can kill *E. faecalis* [43], a pathogen highly resistant to endodontic disinfectants and frequently found in persistent root canal infections [23, 24]. Our research showed that a copper-loaded chitosan-PP scaffold could inhibit the growth of E. faecalis bacteria. This is consistent with a previous *in vitro* study that has also demonstrated a potent antibacterial effect of a chitosan-copper composite against E. faecalis bacteria [44]. However, the antimicrobial activity of this scaffold against other bacteria in infected root canals needs to be studied in further experiments.

Pure chitosan scaffold exhibited a slower wicking rate and degradation rate compared to Chi-PP and Chi-PP-Cu1 scaffolds. The water uptake capacity of a material depends on its hydrophilicity and microstructure [39]. Since chitosan contains NH_2_ and OH groups, it is naturally hydrophilic. However, pure chitosan scaffolds have a lower vertical wicking capacity compared to Chi-PP scaffolds. This could be due to the decreased open porosity of pure chitosan scaffolds. Based on literature, chitosan with a high degree of deacetylation (more than 73%) and high molecular weight shows limited degradation [45], [46]. The medium molecular weight chitosan with ≥ 90% degree of deacetylation used in this study would explain the slow degradation observed for the pure chitosan scaffold compared to Chi-PP and Chi-PP-Cu1. The weight loss of scaffolds containing PP was higher, likely due to the fast degradation rate of PP. Previous reports have shown that PP samples experience a fast degradation rate, with a 50% weight loss within 7 days [47]. In Chi-PP-Cu1, the increased weight loss may be due to the attack at the interface between the polymer and copper, which causes copper and weakly bonded polymeric chains to leach into the solution and degrade the scaffold more quickly. The rate at which scaffolds degrade is also affected by their microstructure, including their porosity content, pore size, distribution, and interconnectivity [48], [49]. Scaffolds that have a higher interconnected porosity tend to degrade faster due to their greater permeability [49]. Although the study did not examine the porosity content and geometry, scaffolds containing PP are highly porous, and the high wicking rate of the PP containing scaffold indicates good interconnectivity between the pores. This could explain why scaffolds containing PP tend to degrade faster than pure chitosan scaffolds.

## 5. Conclusion

This study demonstrates that a novel copper-loaded chitosan-polyphosphate scaffold combines biocompatibility, wicking ability, and biodegradability, while providing short-term infection control and inhibiting calcification. Previous research has mainly concentrated on developing biocompatible scaffolds that can regenerate vascularized pulp tissue for use in regenerative endodontic procedures. However, these studies did not take into consideration the impact of infection and antimicrobial agents on the outcome. This study has successfully created a scaffold that not only has biocompatibility, antibacterial properties, and wicking ability but also is biodegradable and inhibits mineralization. These properties make this copper-loaded Chi-PP an excellent antibiotic-free candidate for an endodontic regenerative scaffold. Therefore, this new approach could have a positive impact in retaining natural dentition on children and adolescents by giving teeth the chance to regain their vitality and continue their growth. Despite the growing interest in developing scaffolds and agents with intrinsic antimicrobial activity to avoid bacterial resistance associated with antibiotic abuse, there have been few promising alternative approaches for scaffold synthesis. Accordingly, this research program will trigger significant advancements in the field of tissue engineering and deliver a new generation of antibiotic free scaffold with optimized antimicrobial properties for endodontic tissue regeneration.

## Funding statement

This study was supported by the Dalhousie Medical Research Foundation and Dalhousie Faculty of Dentistry Research Fund.

## Declaration of Competing Interest

The authors declare that they have no competing interests.

## Notes

### Competing Interest Statement

The authors have declared no competing interest.

