## Supplementary Figures 1 and 2 for "Chitosan-Polyphosphate Scaffold Loaded with Copper for Endodontic Regeneration: A Laboratory Study"

**RATIONALE/JUSTIFICATION (what the investigation contributes to the literature)**

Regenerative endodontics procedures show promise in treating immature teeth with necrotic pulp and apical periodontitis. This procedure involves the replacement of damaged and infected pulp tissue with viable tissue that restores the normal tooth structure and function. Antimicrobials are currently used to control the infection; however, they are cytotoxic to stem cells of the apical papilla (SCAP) and can lead to root canal calcification. Management of these teeth requires a scaffold that can control root canal infection, wick the blood into the canal, and support the viability and differentiation of SCAP while inhibiting intracanal calcification.

**AIM/HYPOTHESIS**

This study aims to develop a composite scaffold made of polyphosphate, a calcium-binding inorganic polymer shown to promote cell proliferation and tissue regeneration, chitosan, a natural antimicrobial polymer that supports stem cell viability and activity, and copper, a metal ion with bactericidal properties.

**ETHICAL APPROVAL (IF APPLICABLE)**

This study received approval from the Nova Scotia Health Research Ethics Board (REB FILE #: 1022150)

**SAMPLES :**

The scaffold was prepared by adding copper (Cu) to chitosan solution, followed by polyphosphate

The stem cell of apical papilla SCAP expressing STRO-1 isolated from immature third molars

**EXPERIMENTAL AND CONTROL GROUPS, INCLUDE INDEPENDENT VARIABLES**

Group 1 – Pure Chitosan (Chi) control group ,

Group 2, and 3 - Chitosan-Copper (Chi-Cu1 and Chi-Cu3) Control group for the effect of PP on copper release,

Group 4– Chitosan-Polyphospate (Chi-PP) Control group for the effect of copper ,

Group 5, 6, and 7 (Chi-Cu1, Chi-Cu2, and Chi-Cu3) Experimental groups .

**OUTCOME(S) ASSESSED, INCLUDE DEPENDENT VARIABLES AND TYPE**

Synthesis of copper loaded Chi-PP scaffold.

Characterization of copper loaded Chi-PP scaffold (Chemical composition and structure).

The ability of this scaffold to support SCAP viability and subsequent odontogenic differentiation and mineralization.

The Antimicrobial activity against *E. feacalis.*

The clinical applicability ( wickability, degradability and the ability to shape in a clinical relevance form).

**METHOD USED TO ASSESS THE OUTCOME (S) AND WHO ASSESSED THE OUTCOME(S)**

Elemental analysis and the structural analysis of various scaffold formulations was assessed by ICP-OES and ATR-FTIR, respecctively. The degrading rate and wicking rate of the scaffold were evaluated over time.

Cell vaiability was evaluated by Alamar Blue assay and a Live/Dead assay, while odontogenic differentiation and mineralization potential were assessed by Immunocytochemistry staining of (DMP-1) and (DSPP) and Alizarin Red staining, respectively.

Antimicrobial property was assessd using colony forming unit analysis. {Assessed by HM}.

**RESULTS**

This study demonstrates that, by combining polyphosphate and chitosan, we could fabricate a scaffold that inhibits bacterial growth by 40 % and supports the viability of fibroblast and SCAP. Adding copper to this scaffold further increased bacterial growth inhibition by up to 68% while preserving cell viability. The immunocytochemistry and Alizarin Red staining revealed that this scaffold also supports the odontogenic differentiation of these stem cells while inhibiting their mineralization potential. Furthermore, this scaffold can be fabricated as a 3D cone-shaped scaffold with a strong vertical wicking ability at a rate of 0.5 mm/s and excellent degradability, with 53 % of the scaffold degraded after 28 days.

**CONCLUSION(S )**

This study demonstrates that copper-loaded chitosan-polyphosphate scaffold combines biocompatibility, wicking ability, and biodegradability, while providing short-term infection control and inhibiting calcification. Therefore, it has great potential as an endodontic regenerative scaffold.

**FUNDING DETAILS**

This study was supported by the Dalhousie Medical Research Foundation and Dalhousie Faculty of Dentistry Research Fund.

**CONFLICT OF INTEREST**

The authors declare that they have no competing interests.

**Supplementary Figure 1:** PRILE 2021 Flowchart

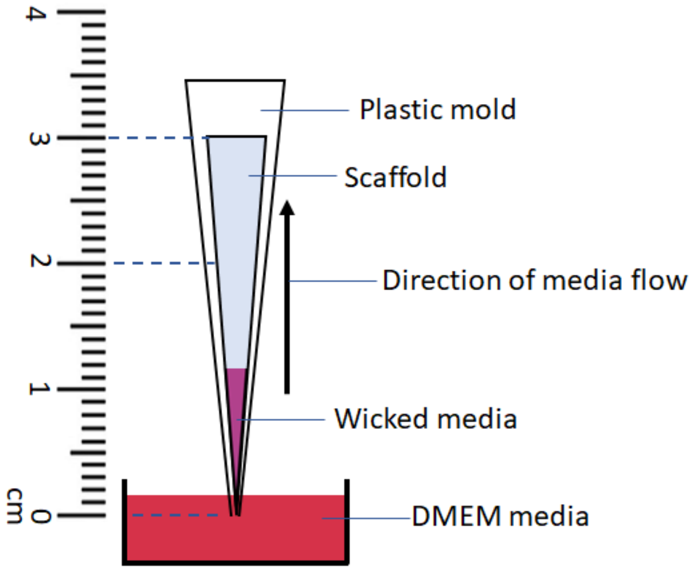


**Supplementary Figure 2:** Schematic of the experimental setup for vertical wicking measurements
